# Treadmill step training promotes corticospinal tract plasticity after incomplete spinal cord injury

**DOI:** 10.1101/2024.10.14.618099

**Authors:** Jaclyn T. Eisdorfer, Hannah D. Nacht, Tess Kowalski, Joshua T. Thackray, Alana M. Martinez, Lance Zymoro, Megan V. Phu, Ridhi R. Hirpara, Burhanuddin S. Danish, Riley Wang, Adyan Khondker, Adam B. Eisdorfer, Max Tischfield, Victoria E. Abraira

## Abstract

Spinal cord injury (SCI) often impairs motor functions such as voluntary movement and fine motor control, with the corticospinal tract (CST) being a crucial pathway affected. While CST-targeted rehabilitation, such as treadmill training, supports motor recovery, gaps remain in understanding the topographical changes within the CST and how they correlate with behavioral outcomes. In this study, we utilized a custom Emx1Cre;LSL-SynGFP mouse line to quantify CST plasticity following moderate contusion SCI, both with and without exercise (treadmill) training. Fluorescent labeling of cortical synapses allowed for detailed visualization of descending CST rewiring, and we assessed its relationship to behavioral outcomes, including kinematics analysis and motivational state. Mice were stratified by motivational state using the Progressive Ratio Assay, and locomotor recovery was evaluated through the Basso Mouse Scale (BMS), joint/limb kinematics, and Motion Sequencing (MoSeq) analysis. Our findings indicate that treadmill training enhances CST rewiring, especially in highly motivated animals, leading to increased synaptic density in the ventral horn and improved BMS subscores. Motivation further influenced specific kinematic parameters, such as toe clearance, while treadmill training significantly improved speed by reducing the stance phase. Results suggest that while treadmill training induces broad beneficial outcomes, motivation may fine-tune recovery, influencing neural circuit and behavioral changes. This suggests multiple mechanisms converge to promote recovery—those we cannot control and those we can. These results underscore the combined role of task-specific training and also perhaps motivation in driving CST plasticity and functional recovery after SCI.

## INTRODUCTION

The corticospinal tract (CST) is crucial for motor functions such as voluntary movement, fine motor control, all of which are often impaired following spinal cord injury (SCI) ^1,2^. Research has consistently supported CST-focused rehabilitation for promoting functional recovery^3,4^, yet significant gaps remain in understanding the topographical changes that occur within the CST post-SCI and how these changes correlate with behavioral outcomes. One promising therapeutic intervention is task-specific training, such as treadmill step training^5^, which has been shown to improve locomotor function after SCI. This approach is believed to promote CST plasticity, facilitating the rewiring of motor circuits necessary for recovery. Furthermore, the motivation to engage in task-specific training has been reported in clinical patients (*personal communication*). The role of motivational state in influencing CST rewiring has been relatively unexplored, despite the fact that motivation is known to modulate motor learning and performance^6^.

Motivational state may thus play a key role in determining the extent of neural rewiring in response to rehabilitation^7^. Highly motivated individuals, or in this case animals, might exhibit more stereotypic and efficient CST connectivity, which could translate into enhanced motor recovery. By examining both treadmill-trained and non-trained mice, while also stratifying them by motivational state, we aimed to uncover the contributions of both training and motivation to CST plasticity and motor function. In this study, we utilized the Emx1Cre;LSL-SynGFP mouse line to label cortical synapses, including those of the CST, enabling us to directly visualize CST rewiring after SCI. We hypothesized that treadmill training would enhance CST plasticity, particularly in highly motivated animals, leading to improved locomotor outcomes. By correlating histological changes in the CST with kinematic outcomes, this study seeks to provide deeper insights into how task-specific training and motivational state together influence functional recovery after SCI. Our findings demonstrate that treadmill training promotes CST rewiring, particularly in the ventral horn, and leads to significant improvements in motor outcomes, as measured by the Basso Mouse Scale (BMS)^8^. Furthermore, we show that motivation affects specific aspects of recovery, such as toe clearance, and may drive the organization of CST connectivity. These results highlight the importance of both training and motivational state in enhancing CST plasticity and motor recovery after SCI.

## METHODS

### Animals

For visualization or manipulation of Emx1Cre, we utilized the Emx1Cre;LSL-SynGFP mouse line. Transgenic mouse strains were used and maintained on a mixed genetic background (CD-1/C57BL/6). Experimental animals used were of both sexes. Mice were provided ad libitum access to food and water and were maintained in a 12-hour light and 12-hour dark cycle within the animal facility at the Division of Life Sciences, Nelson Biology Laboratories, Rutgers University. Spinal cord injury (SCI) surgeries were conducted on these mice at an age of 10 weeks. All housing, surgical procedures, behavioral experiments, and euthanasia were carried out in strict adherence to Rutgers University’s Institutional Animal Care and Use Committee (IACUC; protocol #: 201702589) guidelines, with the experiments following the ARRIVE guidelines.

### Operant conditioning

The Progressive Ratio Assay was conducted with methods previously described^9^ in Med Associates operant chambers (ENV-307W-CT) with MED-PC V and Multi Camera 4K software run on Windows 11. The assay was conducted over 17d, with 8d of reward acclimation in home cages (reward = chocolate pellets), 1d of operant chamber acclimation, 6d of fixed reinforcement training (FR1), and 2d of progressive ratio training (PR2). In the reward acclimation stage, the following was employed: Day 1: Mice were single housed with enrichment, to minimize social interaction differences in home cages outside of the testing; Day 1-3: Mice are weighed for the first three days of acclimation to establish baseline weights (calculated as average weight over three days with no food or water restriction). Along with normal chow, mice are also given 2g of sucrose pellets per mouse (20mg chocolate flavor, Bio-Serv dustless precision pellets) in a petri dish in home cage to familiarize them to the reward; Days 4-5: Mice begin food restriction (1g of chow) + given 4g of sucrose pellets per day; Day 6-8: Mice are still food restricted and only given 4g, 3g, 2g, of sucrose pellets per day respectively. On the chamber acclimation day, mice were brought to the testing room with the Med Associates operant chambers. On the FR1 days: mice were brought to the testing room with Med Associates operant chambers and were awarded 1 sucrose pellet in the food magazine for each nose poke, maxing out at 30 rewards or 90min elapsing. All mice were maintained over 85% baseline body weight. Mice were tested for motivational state during PR2: the first nose poke yields 1 sucrose pellet in the food magazine (instant reward, no delay). Nose poke requirement then scaled cumulatively such that each reward requires 2 more nosepokes than the last (1, 3, 5, 7, 9, etc…). PR2 lasted for 60min.

### Surgical procedures

T9 moderate contusion SCI was following protocols previously described previously^8,10^. Briefly, 10-week-old mice were anesthetized with 5% isoflurane (NDC 66794-017-25, Piramal Critical Care, Inc., Bethlehem, PA, USA), supplemented with 2% during surgery. To sustain body temperature throughtout the duration of the surgery, the animal was placed on a thermo-regulated heating pad. To expose the back for incision, the skin was shaved and sanitized with Betadine (NDC 67618-151-32, Purdue Products L.P., Stamford, CT, USA) and 70% ethanol. Bupivacaine (NDC 0409-1163-18, Hospira, Lake Forest, IL, USA) at 0.1 ml of 0.125% was subcutaneously injected at the incision site. The slow-releasing analgesic, Ethiqa XR (Fidelis Pharmaceuticals, LLC, North Brunswick, NJ, USA) was subcutaneously administered at 3.25 mg/kg. Lubricant ophthalmic ointment (Artificial Tears, NDC 59399-162-35, Akorn Animal Health, Lake Forest, IL, USA) was applied to the eyes following reflexing (pinch) testing. On the skin of the back, a 3 cm incision was made to expose the vertebrae. A laminectomy was performed at T9. Using the IH Infinite Horizon Impactor (Precision Systems and Instrumentation, LLC; model IH-0400), a 50 kDyn moderate contusion was performed. The incision was then closed with wound clips (Reflex-9, 9 mm, Cat. No. 201-1000, Cell Point Scientific, Inc.,Gaithersburg, MD, USA).

### Treadmill training

Treadmill training consisted of locomotion 5d/wk at the following 5 speeds: 5, 10, 15, and cm/s. Training sessions were ∼25 min long and included locomotion at each speed for 4 min, interleaved with 1 min recoveries between speeds. Training took place on a 5-lane treadmill with plexiglass separated lanes. Training concluded after 6 weeks post-injury.

### Behavior

#### Basso mouse scale (BMS)

Locomotor recovery was assessed using BMS^8^. Mice were placed in the center of a small wading pool (open field) with a 44-inch diameter for 4 minutes. Hindlimb movements were scored by two trained observers. The scale is from 0-9, with 0 indicating no movement and 9 representing normality. The lower score was selected in cases of discrepancy between scorers. The subscore, from 0-11 with 0 being no movement and 11 being normality, was also evaluated.

#### Kinematics analysis

Data acquisition was conducted with a custom color four-camera system at a frame rate of 250 Hz while animals locomoted across a custom 5ft horizontal long platform. Each video was calibrated using a ChArUco Board^11^. Steady state locomotion was captured for at least 12-15 steps (required by power analysis). With DeepLabCut (DLC) methods previously described^12–14^, we estimated the locations of the ASIS, hip joint, knee joint, ankle joint, MTP joint, tip of the toe, base of the tail, and tip of the tail. Hindlimb and tail landmarks were manually tracked in ∼1000 frames. The ResNet-50-based model used 95% of the frames for training^15,16^. The effectiveness of joint estimation was determined with a p-cutoff of 0.9. We employed conventional definitions of stride cycles, consisting of swing and stance phases^17^.

### Immunohistochemistry

Mice were deeply anesthetized with 5% isoflurane (NDC 66794-017-25, Piramal Critical Care, Inc., Bethlehem, PA, USA), followed by transcardial perfusion with saline containing 10 U/mL heparin (Cat No. H3393-100KU, Sigma-Aldrich, St. Louis, MO, USA), and 4% paraformaldehyde (Cat No. 15714-S, Electron Microscopy Sciences, Hatfield, PA, USA). Spinal cord columns and brains were dissected. The ventral side of the spinal cord was exposed by removing the ventral bone. Postfixation was done in 4% paraformaldehyde overnight at 4°C. The following day, the tissue was washed 3 times with phosphate-buffered saline (PBS), followed by storage in PBS containing 0.01% sodium azide for further processing.

Injury quantification was performed by exciting 1 cm of each spinal cord containing the SCI site. The excised tissue was cryoprotected with immersion in PBS solution with increasing sucrose concentrations (10%, 15%, 20%) in 4°C for 24h each. The tissue was then incubated overnight with 50% Optimal Freezing Temperature (OCT) (Tissue-Tek, Cat No. 25608-930, VWR, Radnor, PA, USA) medium and 50% 20%-sucrose dissolved in PBS. Serial sagittal cryosections with 20 µm thickness were mounted on Diamond® White Glass Charged Microscope Slides (Globe Scientific Inc, Cat. No. 1358W), and stored at −20°C for immunostaining. Prior to statining, slides were removed from the freezer for 1 hour and left in darkness at room temperature. The tissue was then: washed three times with PBS; blocked with 3% bovine serum albumin (BSA) (Cat No. RLBSA50, IPO Supply Center, Piscataway, NJ, USA), 10% Normal Goat Serum (NGS) (Cat No. 01-6201, Fisher Scientific, Waltham, MA, USA), and 0.3% Triton-X 100 (X100-100ml, Sigma-Aldrich, St. Louis, MO, USA) for 1 hour at room temperature; and stained overnight at 4°C with a primary chicken antibody against the astrocyte marker Glial Fibrillary Acidic Protein (GFAP) (1:1000 in blocking solution; Cat No. 200-901-D60, VWR, Radnor, PA, USA) and a primary rat antibody against myelin basic protein (MBP) (1:50 in blocking solution; Cat. No. MAB386, Sigma-Aldrich, St. Louis, MO, USA). The following day, the tissue was: Washed in five 5-minute washes at room temperature with PBS; incubated for 2 hours at room temperature with corresponding Alexa Fluor 488 (1:500, anti-Chicken coupled to Alexa Fluor 488, Cat No. A11039, Fisher Scientific, Waltham, MA, USA) and Alexa Fluor 546 (1:500, anti-rat coupled to Alexa Fluor 546, Cat No. A11081, Fisher Scientific, Waltham, MA, USA) secondary antibodies diluted in blocking solution; washed five times for 5 minutes at room temperature with PBS; and mounted with glass coverslips (Cat No. 48393-195, VWR) with Fluoromount-G (Cat No. 100241-874, VWR, Radnor, PA, USA).

Spinal cords below the injury site were excised and transverse sections (50 um) were cut using a Leica Vibratome (Mannheim, Germany). Free floating sections were washed in 50% ethanol for 30 min at room temperature and three times in a high salt PBS buffer (HS-PBS) for 10 min each at room temperature. Tissue was incubated in the following primary antibodies made in HS-PBS and 0.3% Triton X-100 (HS-PBST) for 48 at 4°C: primary Guinea Pig antibody against Parvalbumin (PV) (1:1000, Cat No. FR105250, Nittobo Medical Co, LTD, JPN) and primary Rabbit antibody against green fluorescent protein (GFP, 1:500, Cat. No, A11122, Thermo Fischer, Waltham, MA, USA). The tissue was then washed 3 times for 10 min each in HS-PBSt and then incubated in the following secondary antibodies made in HS-PBSt for 24 hr at 4°C: Alexa Fluor 546 (1:500, anti-guinea pig coupled to Alexa Fluor 546, Cat No. A11074, Fisher Scientific, Waltham, MA, USA) and Alexa Fluor 488 (1:500, anti-rabbit coupled to Alexa Fluor 488, Cat No. A11034, Fisher Scientific, Waltham, MA, USA). The following day, tissue was moused on glass slides (41351253, Worldwide Medical Products) and glass coverslipped (Cat No. 48393-195, VWR) with Fluoromount-G (Cat No. 100241-874, VWR, Radnor, PA, USA).

Images of injury sites were acquired with the automated microscope IN Cell Analyzer 6000 (GE Healthcare, Chicago, IL, USA). Four microscope slides were loaded into the system, and the image capturing protocol from the manufacturer was implemented. The tissue sections were precisely located via preview, and high-resolution images were acquired for subsequent analysis. Spinal cord images were acquired using a Leica TCS SP8 tauSTED 3X (Mannheim, Germany) and the accompanying LASX software. CST density and puncta counts were conducted with a custom program using ImageJ.

### Statistical analysis

Kinematics were analyzed using linear mixed effects (LME) models (with the nlme package in R^18^) with fixed effects (injury, time point) and a random effect (mouse). LME models are models well-suited to handling hierarchical, repeated measures data, and data sets that may contain missing or unequal cases, which are typically encountered in SCI studies. Following modeling, two-way repeated measures ANOVA with Wilcoxon rank sum posthoc test was used to determine differences between groups. Differences in density and puncta were calculated with two-way ANOVA with Tukey multiple comparisons posthoc test.

## RESULTS

To investigate how treadmill step training influences corticospinal tract (CST) plasticity after incomplete spinal cord injury (SCI), we utilized the Emx1Cre;LSL-SynGFP mouse line. This mouse line was generated by breeding Emx1Cre males with LSL-SynGFP females, producing offspring in which cortical synapses, including those of the CST, express fluorescently labeled synaptophysin (SynGFP), allowing for the visualization of synaptic rewiring. Following a moderate contusion SCI, we stratified the animals by motivational state using operant conditioning (**Fig 1**). This approach allowed us to detect high-versus low-motivated animals, providing an opportunity to assess how motivation influences CST plasticity (**Fig 1b**). Mice with higher motivation were hypothesized to show enhanced synaptic plasticity in response to treadmill training. All animals subsequently underwent treadmill training, a known promoter of functional recovery after SCI. Using this design, we examined the relationship between CST rewiring and kinematic outcomes while considering the contribution of motivational state. Synaptic rewiring was quantified through fluorescent imaging of SynGFP-labeled CST projections, and motor behavior was evaluated via high-resolution gait analysis.

**Figure 1.**
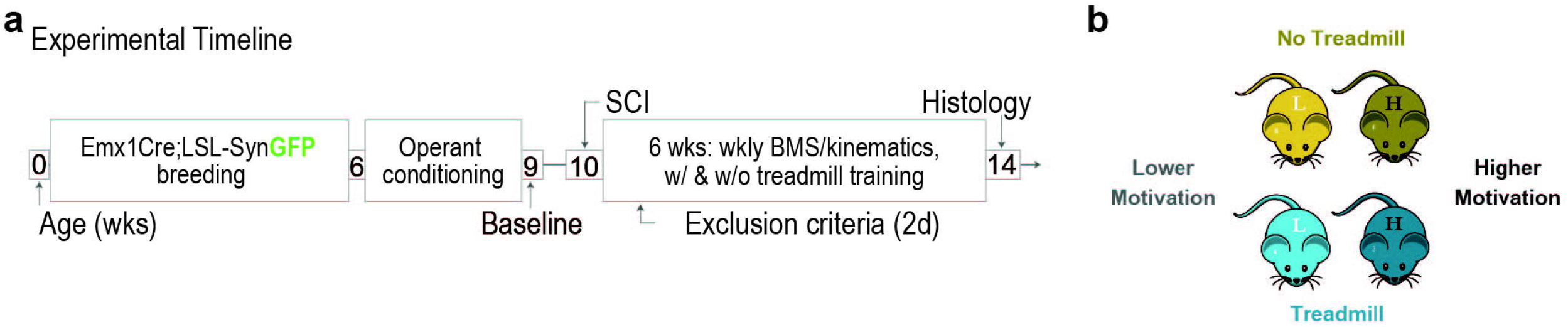
Experimental design. (a) Timeline. Emx1Cre;LSL-SynGFP mice were bred to visualize cortical rewiring, including changes in the CST. Motivational state was stratified using operant conditions (progressive ratio assay). Motivation levels are stratified into “low” and “high” motivators. Weekly BMS and kinematics analysis was performed on treadmill training and no training groups. (b) This study employed four groups: high and low motivated animals that did not exercise and those that treadmill trained.

To assess the impact of treadmill training on corticospinal tract (CST) plasticity and how this may be influenced by motivational state, we quantified CST rewiring in both trained and untrained animals, further stratified by high and low motivation. Treadmill training in highly motivated animals led to an increase in CST puncta onto premotor neurons, specifically deep dorsal horn parvalbumin-expressing interneurons (dPVs) located in laminae IV-V (**Fig 2a**). These inhibitory interneurons play a critical role in corrective motor movements, as previously reported^19^. We observed that the entropy of CST rewiring onto premotor networks in the lumbar spinal cord may be influenced by the animal’s motivational state (**Fig 2b**). High motivation appeared to promote more stereotypic CST connectivity onto premotor networks, whereas low motivation resulted in more entropic, disorganized patterns of connectivity. Additionally, we found that treadmill training significantly increased CST puncta density in the ventral horn (**Fig 2cd**). Using GFP-fluorescing synaptophysin to visualize CST presynaptic terminals in laminae VII-IX, we observed a marked increase in the number of CST terminals in the treadmill-trained groups compared to untrained groups. This increase in CST puncta was particularly evident in highly motivated animals, further suggesting that motivation and treadmill training interact to enhance CST connectivity onto key motor networks involved in locomotor recovery.

**Figure 2.**
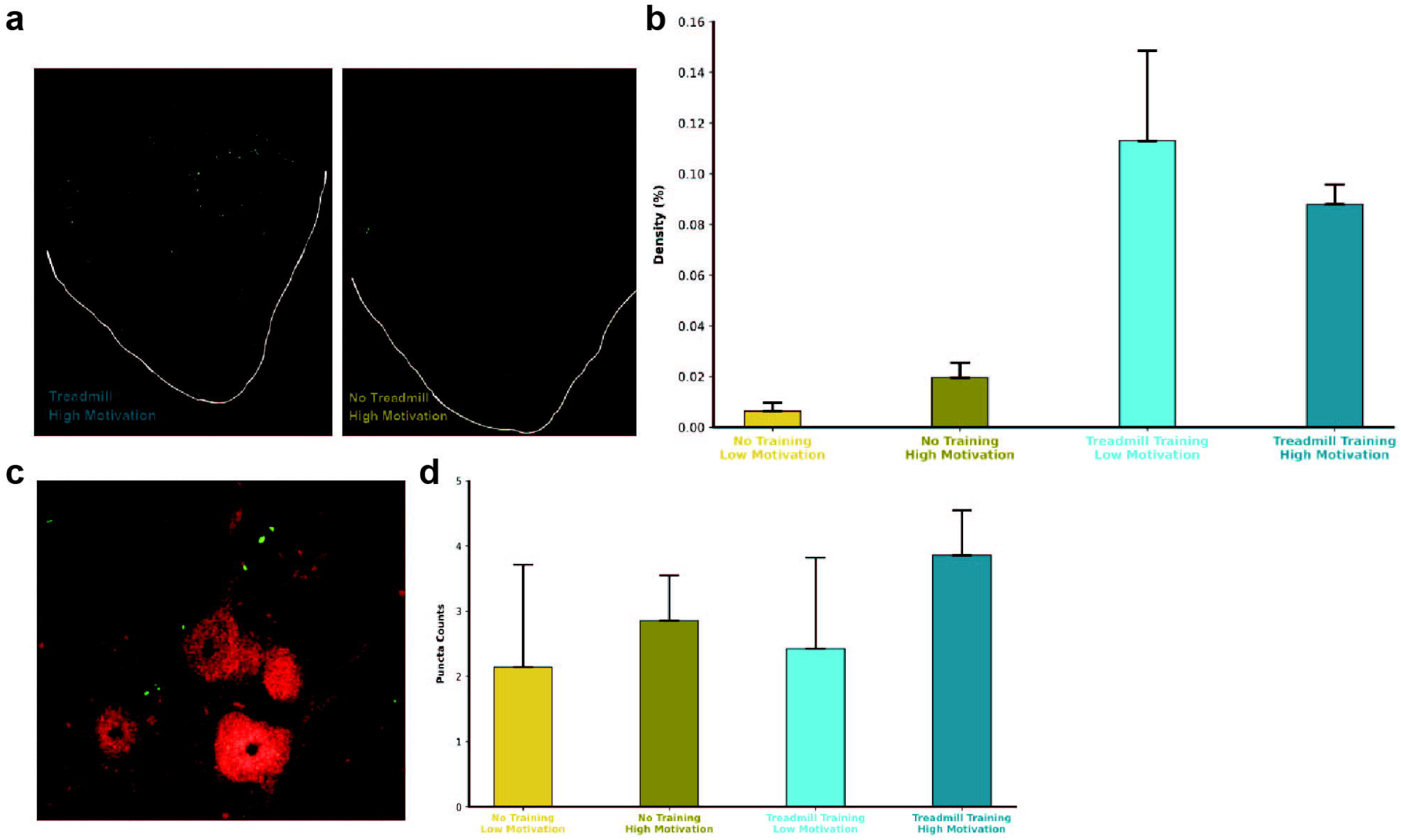
Treadmill training promotes increased synaptic plasticity in the corticospinal tract. (a) Histological image of ventral horn of lumbar spinal cord. (b) Quantification of density in the ventral horn of lumbar spinal cord shows that treadmill training increases CST puncta density in the ventral horn. (c) Histological image of CST puncta onto deep dorsal horn parvalbumin-expressing interneurons (dPVs), inhibitory premotor neurons located in laminae IV-V that are involved in corrective movements. (d) Quantification of (c). Treadmill training with high motivation may also result in increased CST puncta onto premotor neurons. High motivation may thus promote stereotypic CST connectivity onto premotor networks, whereas low motivation may result in more entropic connectivity.

We next examined the kinematic differences between treadmill-trained and non-trained animals, further stratified by their motivational state. These outcomes were assessed using BMS and detailed gait analysis. Kinematics was evaluated while mice walked across a 5ft long horizontal platform which enable capture of individually chosen speed^20,21^. Treadmill-trained mice demonstrated significantly higher BMS subscores compared to non-trained animals, indicating improved overall locomotor recovery post-SCI. The subscores specifically reflected enhanced performance in locomotor functions such as paw and tail position, coordination, and weight-bearing ability (**Fig 3a**). Motivation also appeared to influence kinematic outcomes. Kinematics analysis was performed using 8 hindlimb landmarks: anterior superior iliac spine (ASIS), hip joint, knee joint, ankle joint, metatarsophalangeal (MTP) joint, tip of the toe, base of the tail, and tip of the tail (**Fig 3b**). Mice identified as highly motivated exhibited larger toe clearance off the ground, irrespective of their training status (**Fig 3c**). In contrast, non-trained animals displayed increased instances of toe dragging, with a higher percentage of toe drags occurring in low-motivation mice. This suggests that motivation may play a critical role in fine-tuning specific motor behaviors, such as foot positioning during locomotion. Interestingly, treadmill training was found to significantly increase walking speed, driven by a reduction in the stance phase of the step cycle (**Fig 3d**). However, this effect seemed to be independent of motivational state, suggesting that training itself may be the key factor in promoting faster, more efficient locomotor patterns.

**Figure 3.**
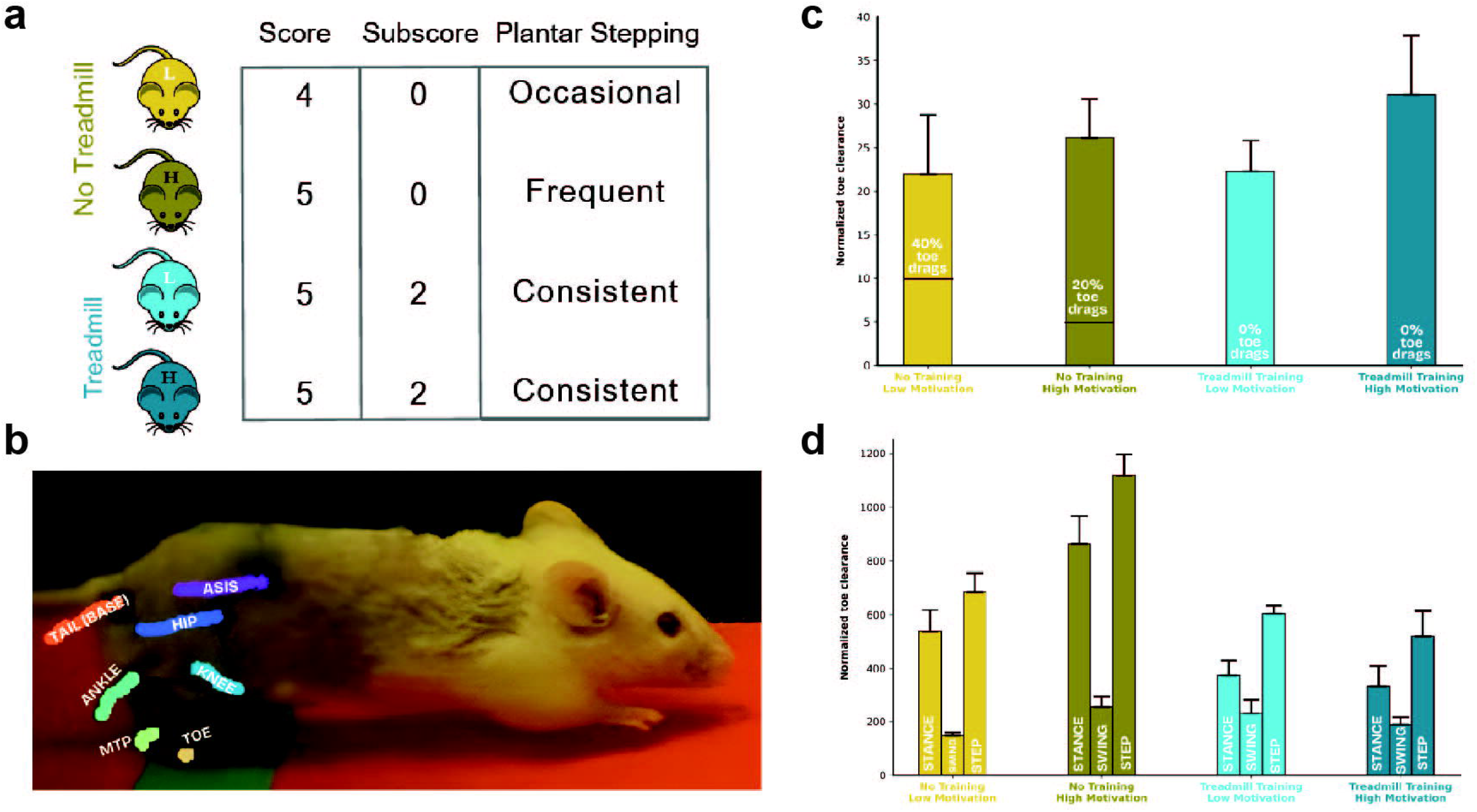
Plastic changes in the CST are correlated to kinematic changes in animals with treadmill training at 21d post-SCI. (a) Treadmill-trained mice exhibit higher BMS subscores at 21d post-SCI. The total BMS score indicates overall locomotive recovery, whereas subscores accumulate scores regarding specific locomotor functions (such as paw and tail position, coordination, etc.). (b) Kinematics analysis included tracking hindlimb landmarks including: ASIS, hip joint, knee joint, ankle joint, MTP joint, tip of the toe, base of the tail, and tip of the tail (not shown). (c) Motivation may drive larger toe clearance at 21d post-SCI. Mice with higher motivation had larger toe clearance off the ground regardless of training or control status. Non-trained animals exhibited toe dragging, with higher percentage of toe drags in those that had lower motivation. (d) Treadmill training increases speed, as dictated by a decrease in the stance phase at 21d post-SCI. Treadmill training, and not motivation, may drive step cycle changes.

## DISCUSSION

Our study investigated the effects of treadmill step training and motivation on corticospinal tract (CST) plasticity and motor recovery following moderate contusion spinal cord injury (SCI). Using the Emx1Cre;LSL-SynGFP mouse line, we visualized CST rewiring and assessed its correlation with kinematic outcomes, revealing the complex interplay between exercise, motivation, and neural plasticity. These findings provide valuable insights into the mechanisms underlying functional recovery post-SCI.

Our results show that treadmill training promoted CST plasticity, particularly in highly motivated animals, as evidenced by increased synaptic density in the ventral horn. This rewiring translated into improved locomotor function, as indicated by higher Basso Mouse Scale (BMS) subscores and enhanced speed due to a reduced stance phase. However, we also observed that motivation alone influenced certain fine motor outcomes, such as larger toe clearance, even in the absence of training. This suggests that motivation may play a role in fine-tuning recovery, influencing both neural circuit dynamics and specific motor behaviors.

Results and gross analysis suggest that while exercise training induces broad, beneficial outcomes, motivation may further refine these effects, operating at both the neural and behavioral levels. If true, this would imply that recovery is driven by multiple converging mechanisms—those we can control through interventions like training, and those we cannot, such as intrinsic motivation. These findings highlight the importance of considering both factors when designing rehabilitation strategies aimed at maximizing recovery.

One limitation of this study is the forced nature of treadmill training, which may not fully capture the impact of voluntary engagement in exercise^5^. Although treadmill training successfully promoted CST plasticity and improved locomotor outcomes, it remains unclear how motivation influences an animal’s willingness to engage in exercise on their own. Future studies will investigate this by examining self-motivated exercise, such as wheel running, to better understand how motivational state affects not only performance but also the decision to participate in rehabilitation.

Overall, our findings demonstrate that task-specific training and motivation interact to shape motor recovery, providing a foundation for future investigations into how intrinsic factors like motivation can be leveraged to optimize rehabilitation outcomes after SCI.

